# Microscaled Cell Surface Proteomics for Cryo-preserved Cells and Tissue Samples

**DOI:** 10.1101/2025.07.18.664488

**Authors:** John R. Thorup, Sarah A. Johnston, Moe Haines, Edwin Sedodo, Kathleen O’Neill, Ronald Drapkin, Namrata D. Udeshi, Michael A. Gillette, Steven A. Carr, Shankha Satpathy

## Abstract

Cell surface proteins (CSPs) regulate key cellular functions and represent valuable targets for diagnostics and therapeutics. Despite advances in proteomic workflows, CSP analysis from cryopreserved or low-input clinical samples remains limited by technical constraints, including reduced membrane integrity, inefficient labeling, and high background. To address these challenges, we optimized and benchmarked two complementary surface enrichment strategies compatible with low-input applications (fewer than 1 million cells) and real-world sample types, including fresh, viably cryopreserved, and dissociated solid tissues.

We systematically compared oxidation-based N-glycoprotein capture and WGA-HRP-mediated proximity labeling across a range of input amounts using both solid tumor (A549) and hematologic cancer (KMS-12-BM) cell lines. The N-glycopeptide method yielded superior specificity in low-input contexts, while WGA-HRP captured complementary CSP subsets. Together, the methods identified more than 700 CSPs, with approximately 175 unique identifications per protocol. Both workflows detected dynamic EGFR internalization following EGF stimulation and maintained high reproducibility (Pearson correlation greater than 0.9) between fresh and cryopreserved preparations.

To extend these findings to tissue-derived samples, we optimized dissociation protocols for healthy endometrium and applied the N-glycopeptide method to cryopreserved dissociated endometrium from three healthy donors. Enzymatic dissociation enabled accurate CSP profiling from fewer than 1 to 2 million cells.

This study provides a systematic comparison of two leading surface proteomics approaches, validates their performance on cryopreserved and low-input specimens, and demonstrates applicability to clinically relevant tissues. Our optimized workflows enable robust surfaceome characterization in translational settings where sample quantity and preservation methods are often limiting, opening new avenues for biomarker discovery and patient stratification.

## INTRODUCTION

Cell surface proteomics is a powerful approach for identifying and characterizing proteins on the plasma membrane (1, 2). These surface proteins play essential roles in cellular communication, signaling, and interactions with the microenvironment, making them key targets for biomarker discovery, drug development, and understanding disease mechanisms.

A range of approaches have been developed for global analysis of the cell surface proteome. The most widely used approaches label surface proteins or glycoproteins with biotin, and the biotinylated proteins are subsequently enriched using streptavidin prior to digestion and analysis by LC-MS/MS. For example, biotin-modified NHS-esters are used to label exposed primary amines like Lys (3, 4). In the widely used method developed by Wollscheid (1, 5–9), cell surface glycoproteins are oxidized with periodate to create functionalizable carbonyls that are reductively aminated with biotinylated reagents. Several proximity-labeling strategies achieve surface-specific biotinylation by tethering promiscuous biotinylators such as HRP or APEX2 to the extracellular membrane. This can be done through glycan recognition, by tethering HRP to the lectin wheat-germ agglutinin, or through non-glycan-targeted anchoring approaches such as DNA-directed localization of APEX2, both of which facilitate labeling of non-glycosylated surface proteins (10). Other approaches for cell surface profiling by MS-based proteomics include incorporation of proximity labeling enzymes into the plasma membrane through engineered mechanisms and the use of membrane impermeable biotin reagents (11–13) to label the local cell surface environment.

As clinical and translational research increasingly relies on archived samples, there is growing demand for workflows compatible with cryopreserved cells and solid tissues. Cryopreservation is a process in which cells and tissues, including patient-derived biopsies, are treated with a cryoprotective agent such as DMSO or glycerol to store biological material for research, diagnostics, and therapeutic development, preserving sample viability for future analysis. While a number of the methods described above have been demonstrated to work not only on fresh cells, but also cells disaggregated from cryopreserved tissue (1, 14), relatively little attention has been given to how cryopreservation itself affects cell recovery and biological integrity. Proteomic analysis of cryopreserved specimens—particularly those derived from patient tissues—presents unique challenges. These samples are often available in limited quantities, necessitating workflows that perform reliably with low input, a historical limitation for surface proteomic methods. In addition, freeze-thaw cycles and tissue dissociation can disrupt membrane integrity, reduce surface labeling efficiency, or introduce degradation artifacts. While multiple studies have shown that dissociation and cryopreservation—individually and in combination—can preserve cell viability, surface marker expression, and biological heterogeneity in single-cell and multi-omic analyses (15, 16), these processes have not been systematically evaluated in the context of surface enrichment proteomics, where intact membrane presentation and efficient labeling are critical for accurate quantification.

To address these challenges, we sought to evaluate and benchmark surface enrichment strategies compatible with both fresh and cryopreserved cell samples with a focus on very low input amounts. Here we systematically compared two orthogonal approaches: (1) peptide capture of labeled N-glycoproteins, a well-established method recently optimized for low-input use (9, 17), and (2) protein-level labeling and enrichment using wheat germ agglutinin conjugated to horseradish peroxidase (WGA-HRP), which has recently emerged as a promising alternative for sensitive surface profiling (10). Both methods were first benchmarked using solid tumor (lung adenocarcinoma) and hematological cancer (multiple myeloma) derived cell lines under controlled conditions, including fresh and cryopreserved preparations, to assess quantitative performance, reproducibility, and sensitivity across a range of input levels.

To evaluate applicability to solid tissue samples, we focused on endometrial tissue, a clinically relevant system with growing interest in surface phenotyping for conditions such as endometriosis (18, 19). As surface proteomic analysis of solid tissues requires viable single-cell suspensions, we optimized dissociation strategies using healthy endometrial tissue to ensure compatibility with downstream labeling and enrichment, and we evaluated the effect of input cells on depth of detection.

## RESULTS

### Comparison of enrichment strategies for cell lines and effect of input amount

To compare performance across input levels, we performed a titration series using 0.1 × 10^6^ to 5 × 10^6^ HEK293T cells with two mechanistically distinct surface enrichment strategies. The glycopeptide-based approach labels surface glycoproteins via periodate oxidation of glycans followed by conjugation to amine-reactive biotin (Figure S1A). After proteolytic digestion, and washing to remove non-glycosylated peptides, N-linked glycopeptides are selectively released by PNGase F. This method is abbreviated here as NGE (N-Glycopeptide Enrichment). Filtering of the data for peptides containing N-X-S/T motifs yields a list of formerly N-glycosylated proteins (Figure 1A) (5). Analyses were performed at the peptide level based on confidently localized deamidation sites in peptides containing the N-linked glycosylation sequence motif retained for downstream analysis (Figure S1B). Some caveats with the approach include the need for the analysis system to be able to robustly and repeatedly detect single or small numbers of formerly glycosylated peptides derived from a protein, and that the ability to detect is dependent on the abundance and capture efficiency of underlying glycopeptides, that can vary depending on glycosylation occupancy and site accessibility. In contrast, the WGA-HRP strategy localizes horseradish peroxidase to cell surface glycoproteins containing NeuNAc (N-acetylneuraminic acid) and/or GlcNAc (N-acetylglucosamine), where it catalyzes oxidation of a biotin-phenol to label proximal tyrosine residues on nearby proteins (Figure S1C) (10, 20, 21). Labeled proteins are then captured and digested directly on-bead, enabling recovery of peptides from both glycosylated and non-glycosylated surface proteins. In contrast to the NGE method, detection in the WGA-HRP is done using non-glycosylated peptides (Figure 1A). A caveat of this method is that it produces a significant amount of intracellular background, complicating confident identification of true cell surface proteins (6). This is also the case for approaches using the Sulfo-NHS-biotin approach (10).

**Figure 1.**
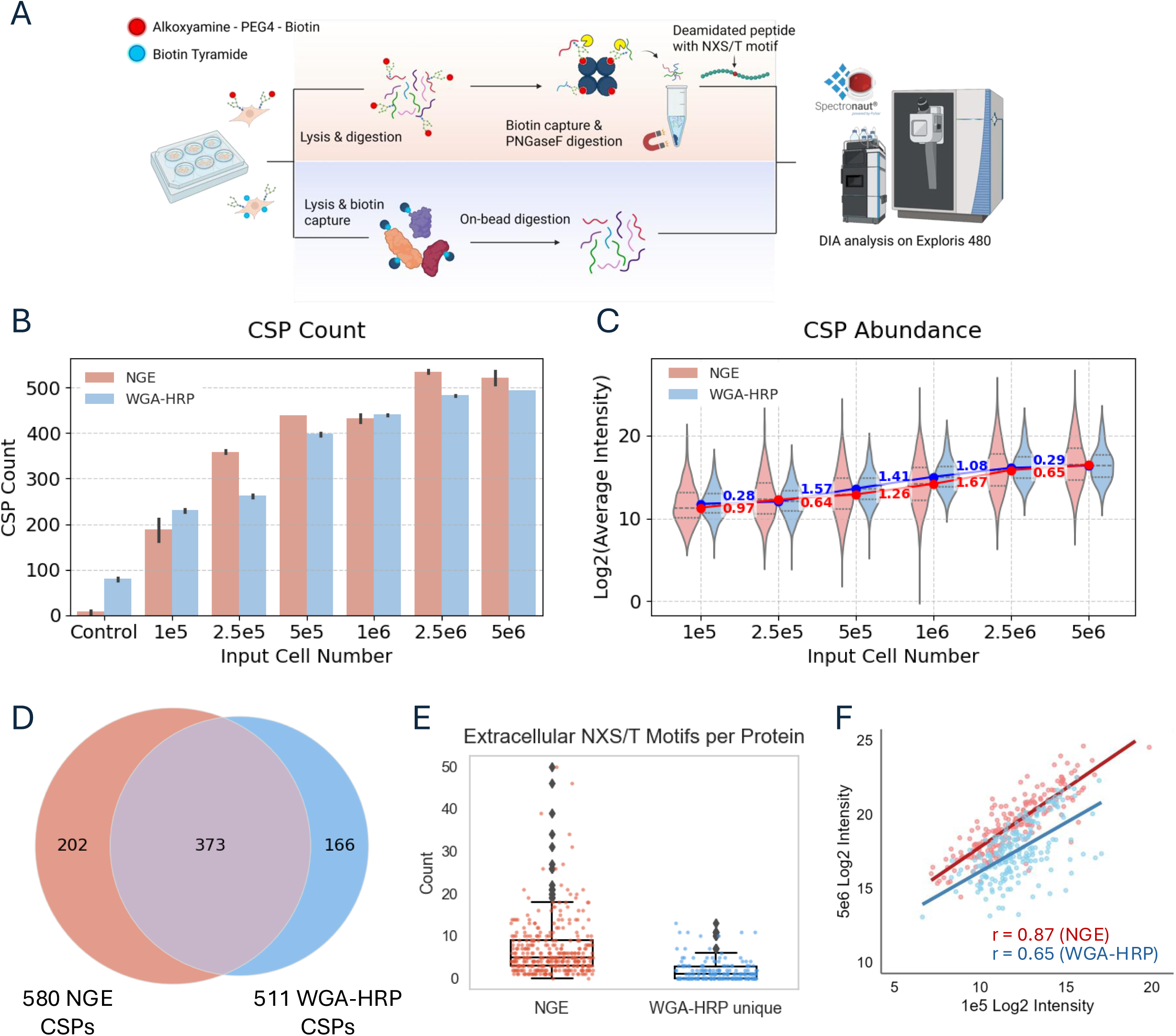
Validation and comparison of glycan oxidation and WGA-HRP labeling protocols across HEK293T cell input titration. **A.** Downstream workflows for cell surface proteomics. For glycan-labeled samples, cells are lysed, proteins digested, biotinylated peptides enriched, and N-glycopeptides released with PNGase F prior to mass spectrometry analysis. For WGA-HRP–labeled samples, cells are lysed, biotinylated proteins enriched, digested on-bead, and resulting peptides are analyzed by mass spectrometry. All data were generated using an Exploris 480 and searched using Spectronaut version 19 (see Methods for details). **B.** Total number of surface protein identifications across input amounts ranging from 1×10^5^ to 5×10^6^ cells for each labeling strategy (N = 3) and average protein intensities. Control samples were generated using 1×10^6^ cells and represent labeling reactions performed in the absence of key reagents: sodium periodate (NaIO_4_) for NGE and WGA-HRP for WGA-HRP labeling. **C.** Violin plots show the distribution of average CSP intensities. Dots represent median log_2_ intensity per input. Lines connect medians across inputs. Numeric labels indicates the log_2_ change in median intensity between adjacent inputs. (i.e., the slope between points) **D.** Venn diagram showing overlap of identified surface proteins across all input amounts between glycan -and WGA-HRP–labeled samples. **E.** Distribution of predicted extracellular NXS/T glycosylation motifs per protein, showing that proteins unique to WGA-HRP have low numbers of extracellular potential glycosylation sites (p-value < 0.01, Wilcoxon rank-sum test). **F.** Correlation of protein intensities quantified in two out of three replicates between low (1 ×10^5^) and high (5×10^6^) input cell amounts.

To benchmark performance in cell lines with a focus on specificity, sensitivity and reproducibility we compared both approaches across an input range of HEK293T cells and analyzed the resulting protein lists using the in silico human surfaceome database (SURFY, Table S1A, (22) to define confident cell surface proteins (CSPs). This provided a consistent framework for evaluating coverage of proteins in the known CSP list, where specificity was defined here as the percentage of proteins identified as CSPs relative to the total number of proteins detected by that method (22). Both enrichment strategies recovered approximately 200 CSPs in the list of known CSPs at the lowest input (1 × 10^5^ cells) and reached a maximum of ∼500 CSPs at 2.5 × 10^6^ cells, with no appreciable gain observed at 5 × 10^6^ (Figure 1B, Table S1B,C). Consistent with these observations, intensity distributions plateaued at higher input levels for both methods, further indicating signal saturation beyond 2.5 × 10^6^ cells (Figure 1C). At intermediate inputs (2.5 × 10^5^ and 5 × 10^5^), the NGE method consistently identified a slightly higher number of CSPs than WGA-HRP (Figure 1B). Cumulatively across all inputs, NGE identified 580 CSPs, while WGA-HRP recovered 511, with an overlap of 373 proteins—highlighting that each method captures a distinct subset of the surface proteome (Figure 1D, Table S1D). Gene ontology analysis showed that proteins uniquely identified by the NGE method were enriched for cell adhesion functions, whereas those unique to WGA-HRP were enriched for ion transport (Figure S1D). Domain-based analysis using topological prediction revealed that many of the proteins uniquely identified by the WGA-HRP method had few N-X-S/T glycosylation motifs in predicted extracellular regions (Figure 1E) highlighting its ability to identify proteins that may not be captured using the NGE method (23). Among these, a subset of ion transporters exhibited structural features, such as short extracellular loops with limited glycosylation, that likely explain their absence from the NGE dataset. A representative topological map of one such protein is shown in Figure S1E (24). Follow-up experiments described below provide further insight into the enrichment of cell adhesion proteins by the NGE approach.

Reproducibility was high for both approaches, with WGA-HRP exhibiting slightly lower CSP coefficients of variation (CVs) across inputs (median CV < 20%). NGE CVs were higher at the lowest inputs but dropped below 20% at 5 × 10^5^ cells and above (Figure S1F). To assess consistency across the input range, we calculated the correlation of CSP quantification between 1 × 10^5^ and 5 × 10^6^ cells for each method. NGE showed higher quantitative agreement across inputs (Pearson r = 0.87), compared to WGA-HRP (r = 0.65) (Figure 1F). This likely reflects greater confidence in low-input NGE measurements, supported by oxidation-minus controls in which CSP identifications dropped to near zero (Figure 1A). In contrast, WGA-HRP identifications in the absence of WGA-HRP remained elevated (∼80 CSPs), consistent with background signal independent of WGA-mediated binding. (Figure 1A). Correlation coefficients between the two enrichment methods at each input level ranged from 0.4 to 0.6, which is unsurprising given the distinct quantification strategies employed—peptide-level quantification from deamidated sites for NGE experiments versus protein level quantification of captured proteins in the WGA-HRP workflow (Figure S1G). Lastly, the proportion of identified proteins classified using Surfy as true CSPs was markedly higher for NGE (∼50%) compared to WGA-HRP (<10%), consistent with the elevated background associated with protein capture–based enrichment strategies (Figure S1H).

### Application to cryopreserved suspension & adherent cell lines

Both strategies were deployed to evaluate both the robustness of low-input workflows across different cell types and their compatibility with cryopreserved material. We used the suspension myeloma cell line KMS-12-BM and the adherent lung adenocarcinoma cell line A549 under fresh and cryopreserved conditions. KMS-12-BM samples were processed at 1 × 10^6^ (low) and 15 × 10^6^ (high) cell inputs for fresh conditions, and at ∼1 × 10^6^ input from cryopreserved material. Approximately 1.3 × 10^6^ cells were frozen using a DMSO-based freezing protocol and stored in liquid nitrogen (see Methods) (Figure 2A). After rapid thawing in warm media and an additional PBS wash, recovered cell numbers based on manual counting were comparable to the fresh low-input condition (Figure S2A).

**Figure 2.**
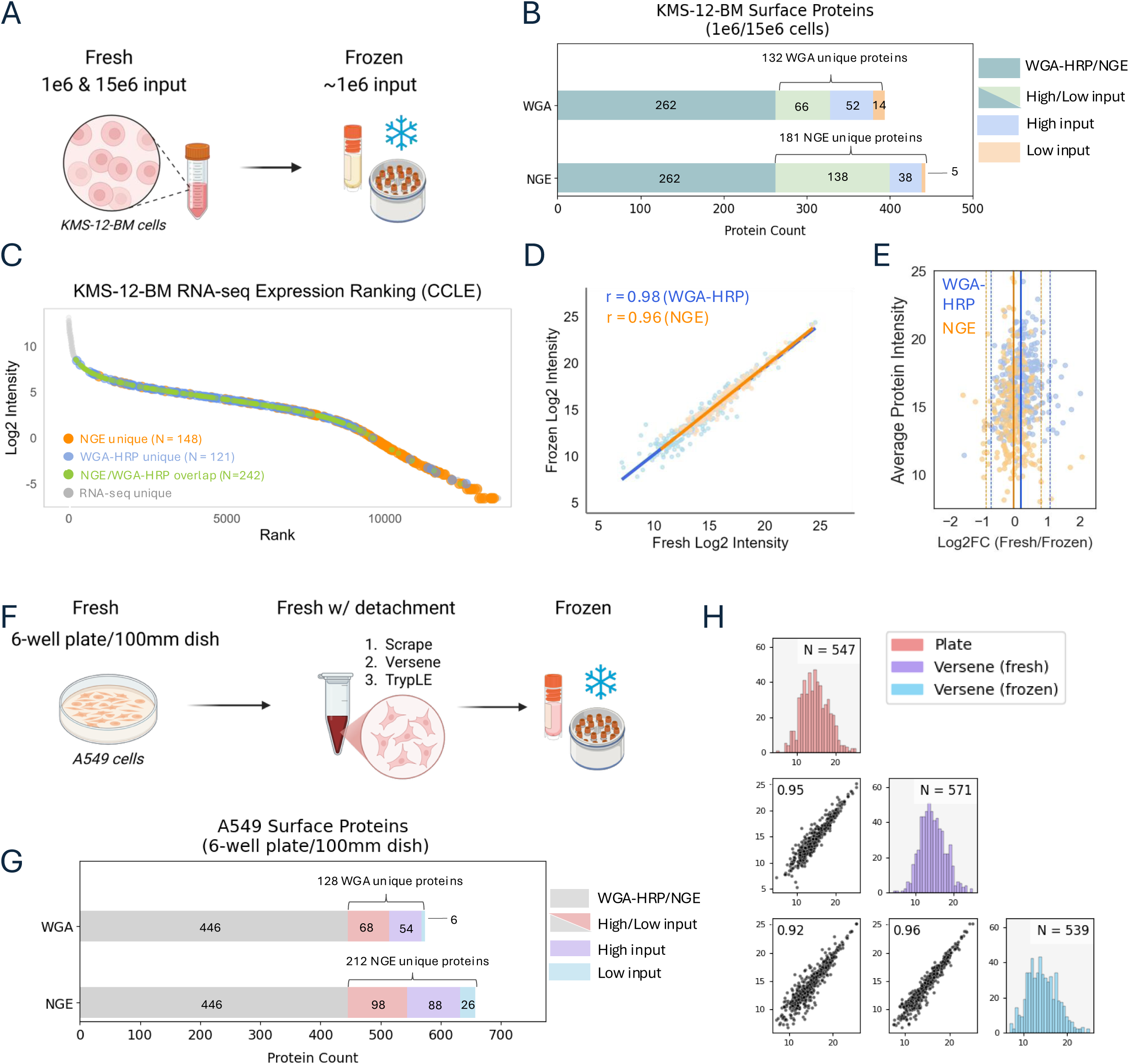
Surface proteomic profiling of cryopreserved and fresh cell lines across input levels. **A.** Experimental design for KMS-12-BM cryopreservation and surface labeling. Cells were frozen in 10% DMSO/FBS, stored in liquid nitrogen, and rapidly thawed for enrichment and analysis. **B.** Bar plot showing CSP overlap between N-glyco and WGA-HRP methods at high and low inputs in KMS-12-BM cells. Partial overlap indicates method-specific subsets. **C.** Rank plot of RNA expression (log_2_ TPM) for surface proteins uniquely identified by NGE vs. WGA-HRP in KMS-12-BM. NGE proteins are enriched at lower transcript levels. **D.** Scatter plots showing strong correlation of CSP quantification between fresh and cryopreserved KMS-12-BM for NGE and WGA-HRP, Pearson r ≥ 0.95 **E.** Log_2_(fresh/frozen) differences are plotted against the average protein intensity across both methods. Solid lines indicate the mean difference (NGE: −0.051, orange; WGA-HRP: 0.171, blue); dashed lines represent ±1.96×SD (NGE: ±0.431; WGA-HRP: ±0.462), capturing the expected spread of variation for each method. **F.** Design comparing A549 surface proteomes across three conditions: fresh adherent, freshly detached, and cryopreserved post-dissociation. Low input samples used 6-well format; high input fresh adherent cells used 100 mm dishes. **G.** Bar plot showing CSP overlap between N-glyco and WGA-HRP at high and low fresh inputs in A549. **H.** Scatter plots and histograms showing consistent surface protein quantification across A549 conditions (plate-based labeling and versene detachment—fresh and frozen) for the NGE method.

Across the fresh input range, we identified approximately 400-440 confident surface proteins (CSPs) per protocol (Figure 2B, Table S2A,S2B). Intra-method overlap of identified CSPs between low and high input levels was high indicating consistent detection across input levels. However, overlap between methods was modest (∼260 CSPs shared), reinforcing the observation that each strategy captures a partially distinct subset of surface proteins (Figure 2B, Table S2C). As seen in HEK293T cells, proteins uniquely identified by WGA-HRP were enriched for ion transport functions, while NGE-specific proteins were dominated by cell adhesion functions with these observed enrichments based at least in part by the presence/absence and abundance of accessible N-glycosylation sites (Figure S2B,C). While the dynamic range of signal intensities appeared similar between methods (Figure S2D), integration with KMS-12-BM RNA-seq data revealed that NGE uniquely captured surface proteins with lower transcript abundance (Figure 2C). These proteins were primarily associated with cell adhesion and likely detected due to the large, heavily glycosylated extracellular domains characteristic of this functional class. A representative topological map of one such cell adhesion protein is shown in Figure S2E.

Comparison of low-input fresh vs. cryopreserved KMS-12-BM samples showed near-complete overlap in CSP identifications for both enrichment methods, with Pearson correlation coefficients ≥ 0.95 for both (Figure 2D, S2F, Table S2D, S2E). To assess the presence of any systematic intensity shift, we performed a Bland-Altman analysis, which confirmed strong agreement, with the vast majority of proteins falling within ±2 standard deviations of the average log_2_ fold change (frozen vs. fresh). Minor shifts in average fold change were observed (–0.051 for NGE, 0.171 for WGA-HRP), reflecting minimal changes in surface signal post-thaw (Figure 2E).

We then extended our analysis to the adherent A549 cell line, which presented the additional challenge of detachment prior to freezing. Cells were labeled in culture dishes at two input scales: 6-well plates (low input) and 10 cm dishes (high input). For the low-input condition, we used the NGE method to compare fresh cells labeled directly on-plate with freshly detached cells using a cell scraper, versene, or trypLE, all of which have been shown to better preserve surface protein integrity compared to classic trypsinization (Figure 2F). Similar to the KMS-12-BM findings, strong overlap was observed between high- and low-input samples within each method, and moderate overlap was observed between the different methods (Figure 2G, Table S2F, S2G). Additionally, RNAseq data from A549 cells validated the trend of NGE’s unique ability to capture a low abundance set of surface proteins (Figure S3A).

In comparing plate-labeled and detached-labeled cells, protein yields following lysis showed that trypLE recovered the largest number of cells (avg. 385 µg), versene and scraping recovered the lowest (avg. 296 µg, 276 µg, respectively), and plate-based labelling recovered an intermediate amount (avg. 328 µg) (Figure S3B). Despite recovering the least total protein, versene-treated and scraped cells recovered the largest number of surface proteins and NXS/T peptides ( Figure S3C,D, Table S2H). While the total number of surface proteins was similar and largely overlapping across methods, differences in NXS/T peptide identification were more pronounced. For example, in versene- and trypLE-treated cells, CSP identifications were 566 and 549, respectively, and NXS/T peptides were 3,461 and 2,915, respectively. The peptides missing in the trypLE condition came from proteins primarily associated with cell adhesion, cell motility, and neurogenesis, suggesting that minor surface shaving occurs during detachment (Figure S3E). Despite these differences, correlation coefficients between all techniques exceeded 0.9, indicating that all are viable options for surface protein detachment while retaining quantitative integrity ( Figure S3F). However, for adherent cell lines, gentle scraping and versene treatment are preferable to trypLE to minimize shaving-related losses. Lastly, we compared frozen A549 cells treated with versene and found a strong correlation (r > 0.9) with both fresh versene-detached and plate-labeled cells (Figure 2H, Table S2H).

### Monitoring dynamic CSP changes using surface proteomics in cell lines

We next evaluated the ability of these surface proteomic workflows to capture biologically meaningful, stimulus-induced changes in surface protein abundance. A549 cells stimulated with EGF, a canonical model of receptor internalization (Figure 3A) (25, 26) was used as a model system to capture dynamic changes and evaluate quantitative changes. This experiment also allowed us to leverage the peptide-level nature of the NGE method for matched surface and global deep-scale proteome analysis. Peptides for global proteome analysis were taken prior to streptavidin enrichment of biotinylated glycopeptides, as non-specific losses during enrichment preclude reliable quantification of flow-through peptides. Although biotinylated glycopeptides themselves are not detectable in the global proteome data, non-glycosylated peptides from surface proteins or intracellular pools of these proteins can still be quantified. In contrast, WGA-HRP captures intact proteins rather than peptides, making it incompatible with paired surface and total proteome analysis from the same input. Following EGF stimulation for 10 minutes or 3 hours in 6-well culture plates, both protocols yielded consistent coverage of A549 cell surface proteins (∼530-560 CSPs) with median CSP CVs ∼20% across biological replicates (N=3). (Figure S4A,B, Table S3A, S3B). Surface EGFR levels, as measured by the NGE workflow, were significantly reduced at both time points, consistent with rapid internalization (Figure 3B). In contrast, global proteome EGFR abundance remained essentially unchanged at 10 minutes, reflecting receptor internalization without degradation, and showed a pronounced decrease by 3 hours, consistent with lysosomal degradation (Figure 3C, Table S3C).

**Figure 3.**
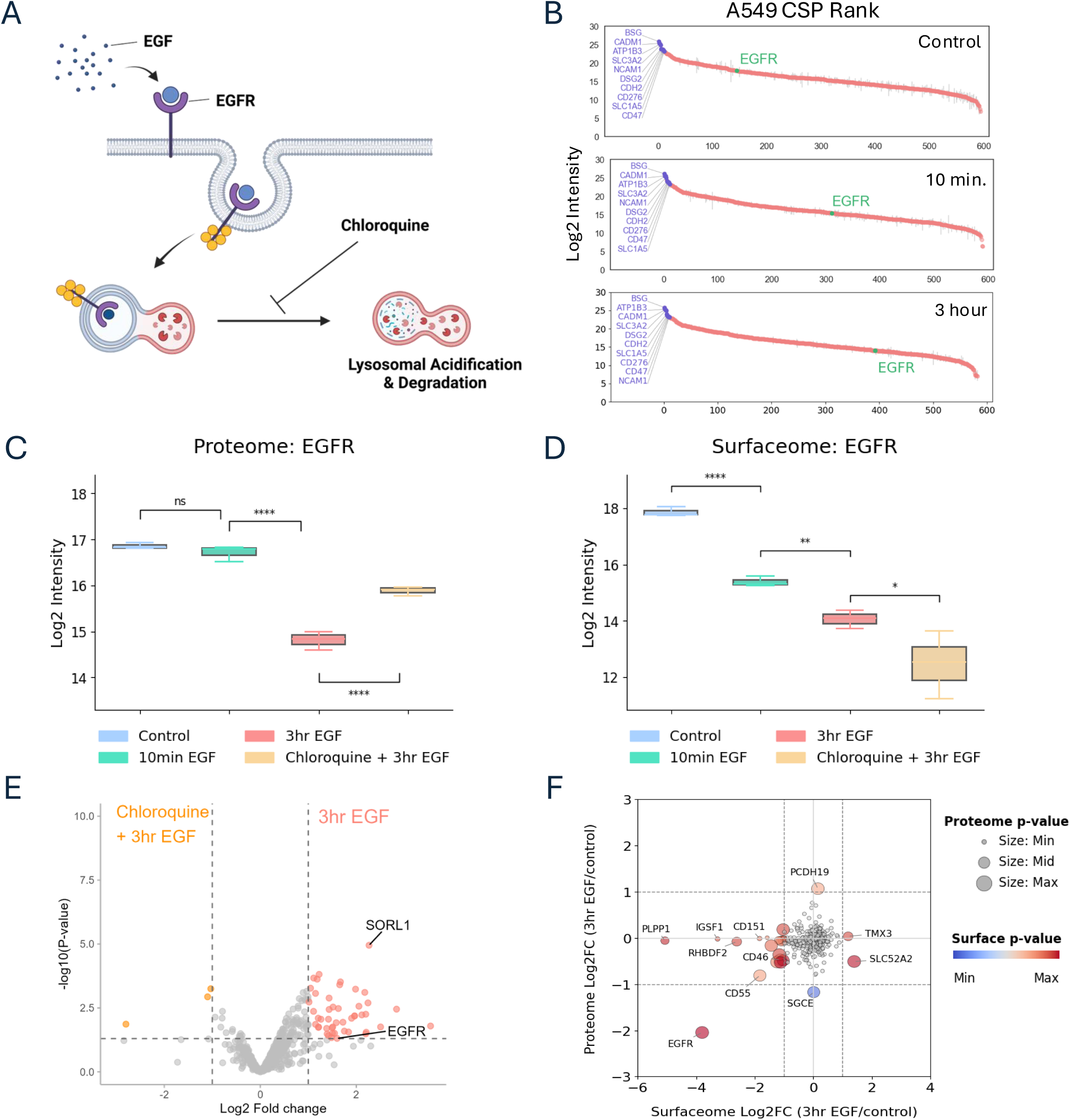
Contrasting surfaceome and proteome dynamics of receptor trafficking in response to EGF stimulation. **A.** Schematic of EGF-induced EGFR internalization and lysosomal degradation, with or without inhibition of lysosomal function by chloroquine. **B.** Rank plot of N-glyco–based surfaceome data from control, 10 min. EGF, and 3 hr. EGF conditions, highlighting EGFR and high-abundance stable surface proteins. **C.** Total EGFR protein levels from N-glyco labeling of A549 cells across four conditions: untreated control, 10-minute EGF, 3-hour EGF, and chloroquine pretreatment followed by 3-hour EGF stimulation. EGFR abundance is quantified from surfaceome data (N = 3). Significance determined by unpaired two -tailed t-tests: *P* > 0.05 (NS), *P* < 0.0001 (****). **D.** Surface EGFR levels from the same conditions and samples, measured by whole-proteome analysis (N = 3). Significance determined by unpaired two-tailed t-tests: *P* < 0.05 (*), *P* < 0.01 (**), *P* < 0.0001 (****). **E.** Volcano plot comparing surfaceome profiles at 3-hour EGF vs. chloroquine + 3-hour EGF, showing reduced surface EGFR and downregulation of proteins involved in EGFR recycling in chloroquine-pretreated samples (log_2_-fold change > |1|; p < 0.05 by t-test). **F.** Comparison of surfaceome and proteome log_2_ fold changes for overlapping proteins for one time point (3 hr. vs. control), highlighting that significant regulation is primarily detected in the surfaceome data (log_2_-fold change > |1| for proteome and surfaceome data).

To test whether receptor internalization and degradation could be blocked, we pretreated cells with chloroquine at levels shown to inhibit autophagy prior to 3-hour EGF stimulation (27). Chloroquine led to increased total EGFR levels, consistent with impaired lysosome mediated-degradation, but resulted in even greater depletion of surface EGFR, suggesting continued internalization without efficient recycling (Figure 3D). When comparing surface profiles between 3-hour EGF-stimulated samples with and without chloroquine pretreatment, we observed a significant reduction in SORL1 alongside EGFR (Figure 3E). SORL1 is known to mediate ERBB2 recycling and was recently implicated in EGFR trafficking (28, 29). These findings suggest that lysosomal disruption may interfere with SORL1 function, leading to impaired recycling and enhanced surface depletion of EGFR. While prior studies have shown that SORL1 silencing can impair lysosomal activity, our data reveal the inverse relationship—highlighting how lysosomal dysfunction can potentially reveal SORL1’s role in trafficking regulation (29).

To further assess whether the CSP changes observed could be obtained just using whole proteome analysis, we compared the log fold-change of CSPs identified in both the surface and total proteomes following 3-hour EGF stimulation. EGFR was the only protein to show consistent downregulation in both the surface and total proteomes, reflecting its well-characterized internalization and degradation dynamics. In contrast, the majority of surface-regulated proteins showed little to no change at the total proteome level, while significant fold changes were observed exclusively in the surface-enriched fraction (Figure 3F). These findings highlight the limitations of global proteomics in capturing spatial and dynamic remodeling and underscore the value of surface-specific enrichment for detecting dynamic regulation.

Using WGA-HRP enrichment, we similarly observed EGFR surface depletion following 10-minute and 3-hour EGF stimulation (Figure S4C). However, the chloroquine-treated condition showed only a modest, non-significant decrease in EGFR alongside a significant reduction in SORL1 (**Figure 4SD,E**). This may reflect labeling and enrichment of EGFR by WGA-HRP in the intracellular pool or compression of the actual fold change due to the significantly higher background protein levels in the WGA-HRP method, both of which are potential limitations in distinguishing surface-localized vs. internal protein pools.

**Figure 4.**
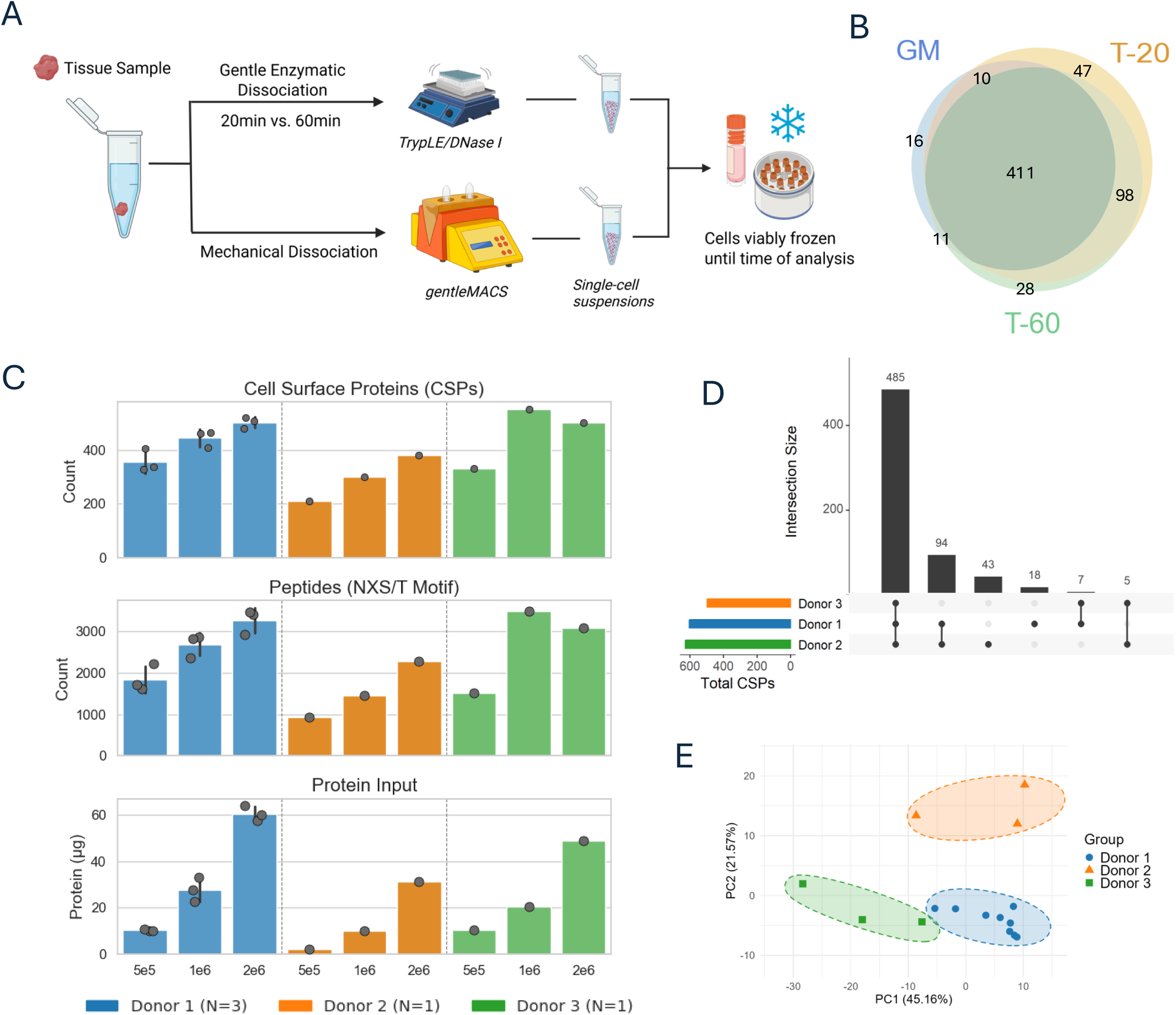
Optimization and evaluation of a low-input surface proteomics workflow using dissociated endometrial tissue. **A.** Schematic of the dissociation and cryopreservation workflow. Fresh endometrial tissue was subjected to either enzymatic digestion with TrypLE/DNase I for 20 minutes (T-20) or 60 minutes (T-60), or mechanical dissociation using the gentleMACS system (GM). Resulting single-cell suspensions were cryopreserved and later used for surface proteomic analysis. **B.** Overlap of confident CSP identifications among the three dissociation protocols. The majority of proteins identified in the GM and T-60 conditions overlapped with those captured using T-20. **C.** Cell surface protein (CSP) counts, NXS/T motif-containing peptide counts, and total protein input for each titration sample using the T-20 method. Inputs of 0.5 × 10^6^, 1 × 10^6^, and 2 × 10^6^ cells were analyzed for three donors, with one donor profiled in triplicate. CSP yields remained robust even at the lowest input levels. **D.** UpSet plot showing the number of CSPs shared and uniquely identified across donor–input combinations. High overlap in CSP identifications was observed across donors and input levels. **E.** Principal component analysis of CSP intensities after experiment-wide median intensity normalization reveals clear donor-specific clustering, demonstrating reproducibility and biological distinction across samples.

### Surface proteomic profiling of cryopreserved single-cell suspensions from solid tissue

To enable surface proteomic profiling of clinical lesions with limited input material, we optimized a dissociation and cryopreservation workflow using healthy human endometrial tissue as a model system. We sought to establish a protocol that maximizes post-thaw viability and surface labeling compatibility while working within input constraints typically encountered in this clinical context. Using endometrial tissue from a single donor, initially, we compared three dissociation protocols (see Methods): (1) gentle enzymatic digestion with TrypLE and DNase I for 20 minutes at 37°C (T-20), (2) same enzymatic digestion for 60 minutes (T-60), and (3) mechanical dissociation using the gentleMACS Dissociator (GM) (Figure 4A). Prior experiments performed in our group suggested that TrypLE has superior ability to preserve cell surface epitopes, critical for downstream proteomic applications for complex solid tissue. Unlike traditional enzyme blends such as Liberase, TrypLE also offers higher purity and specificity, minimizing degradation of surface markers during dissociation. Tissue was manually divided into 150-200 mg wet weight aliquots and processed in replicate to generate technical pairs. Cell yields ranged from 20,000– 30,000 cells per mg of tissue across all conditions, resulting in total recoveries sufficient to generate cryopreserved aliquots containing >1.5 × 10^6^ cells (Figure S5A).

After thawing, cell viability remained high for both enzymatic conditions, with ∼70% of input cells recovered, whereas the gentleMACS protocol yielded substantially lower post-thaw recovery (∼40%) (Figure S5B). This reduced viability may reflect accumulated mechanical stress during dissociation, which can compromise membrane integrity or increase sensitivity to freeze–thaw damage. Surface proteomic analysis on the cryopreserved replicates yielded an average of approximately 400 confident surface protein (CSP) identifications (Figure S5C, Table S4A). CSP profiles were highly consistent across T-20 and T-60 replicates, indicating minimal impact of extended incubation time, and nearly all proteins identified in the gentleMACS condition overlapped with those captured by the enzymatic protocols (Figure 4B). Based on comparable proteomic depth, high post-thaw viability, and reduced incubation time, the T20 protocol was selected for subsequent use.

To evaluate the performance of our optimized workflow under clinically relevant constraints, we selected the T-20 protocol for a titration series using cryopreserved cells from three independent endometrial donors. Cell inputs of 0.5 × 10^6^, 1 × 10^6^, and 2 × 10^6^ were chosen to reflect the range of yields that might be obtained from dissociated endometriotic lesions. Each donor contributed a full titration series; for one donor, triplicate input sets were generated from independent pieces of tissue. Tissue input weights for dissociation ranged from 158–170 mg, yielding higher cell recoveries than in the optimization experiments, at approximately 60,000–90,000 cells per mg of tissue (Figure S5D). Across all conditions, surface proteomic analysis yielded high CSP identification rates (typically >400 CSPs per sample), with only modest reductions observed at the lowest input level (Figure S4C, Table S4B). Protein identifications demonstrated strong overlap between input amounts, indicating consistent enrichment performance even with limited material (Figure S4D). Principal component analysis further showed clear donor-specific separation, emphasizing both the reproducibility of the method and its sensitivity to underlying biological variation (Figure 4E). These results establish the feasibility of robust surfaceome profiling from limited clinical material.

## DISCUSSION

This work addresses two key gaps in the field: first, by providing a direct, head-to-head comparison of two mechanistically distinct, low-input-compatible surface enrichment strategies under standardized conditions; and second, by demonstrating that these optimized workflows can be successfully applied to cryopreserved cell lines and single-cell suspensions derived from solid tissues. Together, these advances broaden the applicability of surface proteomics to clinically relevant, low-input samples and support its use in translational research settings.

Despite growing interest in surface proteomics for translational applications, there remains a lack of standardized frameworks for low-input or cryopreserved samples. By systematically evaluating two distinct enrichment strategies across input levels, preservation states, and dissociation protocols, this work provides practical guidance for implementing surfaceome profiling in resource-limited or clinically derived contexts. While various protein capture methods employing different chemistries have been explored individually (10), this study offers the first direct comparison of two mechanistically distinct surface enrichment approaches—from labeling strategy through downstream enrichment. Both strategies use commercially available reagents and are straightforward to implement, enabling high-quality surfaceome coverage from inputs below 1 × 10^6^ cells—cell numbers that previously would have been considered insufficient for deep profiling. Additionally, all analyses were performed on a single Exploris 480 instrument, using manual implementations of the protocols. While others have explored automation to improve throughput, our results demonstrate that robust low-input surface profiling can be achieved with accessible instrumentation and hands-on workflows (9, 17). We anticipate that this comparison will help researchers select appropriate methods based on sample characteristics, experimental goals, and analytical constraints.

In our optimizations and comparative analyses, we leveraged SURFY as a uniform way to assess the degree of cell surface protein coverage and selectivity of the method for CSPs vs background proteins. However, use of these curated lists limits the ability to detect novel, previously unknown true cell surface localized proteins. Including proteins with predicted transmembrane domains, signal peptides, or surface annotations from databases like UniProt—will likely yield broader coverage, and may improve the ability to detect novel CSPs.

We selected the NGE approach for tissue-derived applications based on several key advantages. It demonstrated high confidence in low-input surface protein quantification, with consistent CSP recovery and strong correlation across input levels. Its reliance on glycan-directed labeling and selective release of N-linked peptides resulted in lower background and reduced need for aggressive post-hoc filtering, enabling more complete analysis of all identified proteins. Additionally, the method showed enhanced dynamic range and successfully captured biologically meaningful changes in surface protein abundance, as seen in our EGF stimulation model. These features make the NGE workflow particularly well suited for complex or limited-input samples such as cryopreserved dissociated tissue. Finally, while we selected the 20-minute TrypLE/DNase I dissociation protocol based on performance and handling efficiency, we acknowledge that optimal dissociation strategies may differ across tissue types and should be empirically evaluated based on cell type and structural composition.

The merits of the WGA-HRP approach should not be overlooked. Protein-level capture may offer improved quantification within a sample due to broader sequence coverage, particularly in cases where differential glycosylation influences the yield and apparent abundance of target proteins, as we observed for certain cell adhesion molecules. Moreover, differences in instrumentation or refinements to labeling workflows may mitigate some of the limitations we encountered related to background and sensitivity. Our control experiments (i.e., oxidation-minus for NGE and WGA-HRP-minus for protein capture) highlighted potential background interference in protein-level enrichment, which may complicate accurate CSP quantification at the lowest input levels and require examination of much larger proteins lists that are dominated by secreted and intracellular background proteins. This is a general challenge with protein capture-based strategies. A recent study employing protein-level capture of glycoproteins using labeling chemistries analogous to those used in our NGE strategy similarly reported background signal as a concern (6).Thus, for applications involving extremely limited material (e.g., <5 × 10^5^ cells), the NGE approach may offer greater reliability for quantitative surface profiling.

Looking ahead, this work expands the applicability of surface proteomics to clinically relevant materials that have traditionally been inaccessible to enrichment-based workflows. In particular, it enables investigation of the surface proteome in contexts such as therapeutic response, receptor trafficking, and immune evasion—where cell surface remodeling can have functional and clinical significance. Ongoing refinement and validation of flexible, low-input-compatible workflows will be critical for realizing the full translational potential of surface proteomics, and the approaches described here contribute toward that goal.

Reliance on curated surfaceome databases like SURFY, while effective for assessing coverage and selectivity of cell surface proteins (CSPs) and background proteins in context of method evaluation and optimization, inherently limits the ability to identify novel or previously unknown true cell surface localized protein. This means that while the study can meticulously validate and quantify known CSPs, filtering may potentially overlook important, yet unannotated, cell surface markers or therapeutic targets. To circumvent this, careful curation will be necessary for the list of proteins that do not overlap with the CSP list. The shorter list generated by NGE makes it easier. A further consideration is that this method relies on the cellular membrane remaining intact during dissociation, thus excluding its application to certain viably preserved tissues, including formalin-fixed samples. Finally, the number of cells per mg of tissue may vary across different tissue types, and pilot experiments are necessary to establish the tissue needed to get at-least a million cells as is required for these experiments.

## DATA AVAILABILITY

Proteomics data generated will be made available upon publication via MassIVE

## SUPPLEMENTAL DATA

Table legends:

**Table S1**: Cell surface proteomics data associated with the HEK Titration and DIA method details

**Table S2**: Cell surface proteomics data associated with KMS12BM and A549 input, cryopreservation, and detachment comparisons

**Table S3**: Cell surface proteomics and global proteomics data associated with EGF-stimulated A549 cell line experiment

**Table S4**: Cell surface proteomics data associated with endometrial tissue dissociation evaluation and input titration

## ACKNOWLEDGMENTS

This work was supported by the following awards and grants - Broad Institute SPARC (#800444) awards to S.S and S.A.C, National Cancer Institute (NCI) Clinical Proteomic Tumor Analysis Consortium (CPTAC) grants U24CA270823 to S.S, M.A.G and S.A.C, U01CA271402 to M.A.G and S.A.C and, Dr Miriam and Sheldon G. Adelson Medical Research Foundation to S.A.C and N.D.U. BioRender was used for a subset of figures.

## AUTHOR CONTRIBUTIONS

J.T. and S.S. conceptualized this study. J.T. executed all cell line and tissue proteomic experiments. M.H. assisted with MS sample preparations. A.J. and E.S. conducted tissue homogenization as depicted in Figure 4, under the guidance of R.D. and K.N. All authors participated in data analysis, data interpretation, and manuscript preparation.

## METHODS

### Experimental Model and Study Participant Details

HEK293T and A549 cells were obtained from internal Broad Institute repositories, and KMS-12-BM cells were provided by the Ghobrial laboratory at Dana-Farber Cancer Institute. KMS-12-BM cells were cultured in RPMI-1640 medium (Gibco) supplemented with 10% fetal bovine serum (FBS), while HEK293T and A549 cells were maintained in DMEM (Gibco) with 10% FBS. All cells were grown at 37°C in a humidified incubator with 5% CO_2_.

Cryopreserved single-cell suspensions from dissociated endometrial tissue were provided by the O’Neill Laboratory and collaborators at the University of Pennsylvania School of Medicine. Tissue dissociation and cryopreservation were performed on-site prior to shipment. Patients were consented {for use}.

### Cell Culture and Reagents

For stimulation experiments, A549 cells were treated with 100 ng/mL epidermal growth factor (EGF; PeproTech) in DMEM containing 2% FBS. For chloroquine pretreatment conditions, cells were first incubated for 2 hours in DMEM with 2% FBS containing 10 µM chloroquine (Selleck Chemicals), after which EGF was added directly to the same medium and cells were stimulated for an additional 3 hours.

For cryopreservation, cells were resuspended in 10% DMSO in FBS, slowly cooled in an ethanol-based freezing container at –80°C overnight, and transferred to liquid nitrogen the following day. After an additional overnight storage in liquid nitrogen, cells were thawed and processed. For suspension cells (KMS-12-BM), fresh and freshly thawed cells were labeled and processed in parallel. For adherent A549 cells, fresh cells were labeled immediately, while the dissociated cells were cryopreserved using the same protocol. Thawed A549 cells were then labeled and both fresh and frozen samples were processed together through all downstream steps.

### Tissue collection

Endometrial biopsies were obtained from the study participants using Pipelle Endometrial Biopsy Curette (3.1 mm Tip, Cooper Surgical). Biopsies were obtained in the luteal phase of the menstrual cycle in ovulatory women using the date of the first day of their last menstrual period to establish the cycle day of biopsy. Tissue was placed in ice-cold calcium and magnesium free Dulbecco’s phosphate-buffered saline (1X PBS), rinsed twice to remove excess mucus and blood and placed on ice for transport back to the laboratory.

### Tissue dissociation

#### T-20 & T-60 Protocols

Tissue was weighed and placed into a petri dish and tissue was finely minced with two razor blades for 30 seconds. Using a large-bore transfer pipette, tissue was collected and placed in ice-cold digestion media comprised of TrypLe Express (Gibco) with DNase I (Roche, #10104159001, 30ug/mL stock concentration, 30ng/mL working concentration). For the T-20 protocol, tissue was incubated in digestion media for 20 minutes at 37°C in a shaking incubator (75 rpm), and a wide-bore transfer pipet was used to mix the tissue every 5-7 minutes during the incubation. For the T-60 protocol, tissue was incubated in digestion media for 60 minutes at 37°C in a rotating incubator (40 rpm), and a wide-bore transfer pipet was used to mix the tissue every 15 minutes during the incubation. Cell suspensions were then applied to a 70-μm cell strainer (Corning, #431750) over a 50 mL conical tube and the suspension was pushed through the filter using the plunger end of a syringe. Digestion media was diluted with an ice-cold MACS buffer (PBS (without calcium, without magnesium), 2mM EDTA, 2% fetal bovine serum) by pouring MACS buffer through the filter. The sample was pelleted at 300g for 5 minutes at 4°C and the supernatant was discarded. The cell pellet was resuspended in 3-5 mL of ACK Lysing buffer (Gibco, #A1049201) and incubated at room temperature for 5 minutes. Ice-cold 1X PBS was added to the cell suspension to stop the lysis reaction, and the suspension was spun at 300g for 5 minutes at 4 °C. The supernatant was discarded and the cell pellet was resuspended in 1X PBS + 0.04% BSA (Sigma, #B2518-100MG). Cell viability was determined by Trypan Blue staining and hand counting with a hemocytometer. Cells were cryopreserved in 90% FBS with 10% DMSO and stored in liquid nitrogen until ready for analysis.

### GM (gentleMACS) Method

Tissue was dissociated according to protocol **<u>dx.doi.org/10.17504/protocols.io.bvy8n7zw.</u>** The final cell pellet was resuspended in 1X PBS + 0.04% BSA (Sigma, #B2518-100MG). Cell viability was determined by Trypan Blue staining and hand counting with a hemocytometer. Cells were cryopreserved in 90% FBS with 10% DMSO and stored in liquid nitrogen until ready for analysis.

### N-glycoprotein Labeling (NGE Method)

For tube-based labeling, cells were transferred to either 0.5 mL or 1.5 mL Protein LoBind tubes (Eppendorf) and washed twice with warm phosphate-buffered saline (PBS). For adherent cell lines, PBS containing calcium and magnesium was used. All subsequent steps were performed on ice unless otherwise noted. Cells were incubated with 1.6 mM sodium metaperiodate (Thermo Fisher Scientific) in PBS (pH 6.5) for 15 minutes at 4°C, followed by a single wash with 1% FBS in PBS (pH 6.5). Surface aldehydes were then labeled with 500 µM alkoxyamine–PEG4–biotin (Thermo Fisher Scientific) supplemented with 5 mM 5-methoxyanthranilic acid (Thermo Fisher Scientific) in 1% FBS/PBS (pH 6.5) for 30 minutes at 4°C. Cells were washed once more with 1% FBS/PBS (pH 6.5) and twice with PBS (pH 6.5). For plate-based labeling (6-well plates and 10 cm dishes), the same protocol was applied, with reagent volumes scaled to fully cover the adherent monolayer and all steps performed using manual aspiration. After the final PBS wash, cells were gently scraped into tubes and processed identically to suspension samples. All centrifugation steps were performed at 650 rpm for 2.5 minutes, and resulting pellets were frozen at –80°C until lysis.

### N-glycoprotein Lysis, Digestion, and Enrichment

Samples were processed similar to previously published methods (9), with minor modifications. Cell pellets were lysed in 8 M urea, 50 mM ammonium bicarbonate (ABC), 2% sodium dodecyl sulfate (SDS), and 1 mM magnesium chloride (MgCl_2_), and mixed at 1,000 rpm for 10 minutes at room temperature using an Eppendorf ThermoMixer. Lysates were treated with 1 µL Benzonase (Millipore Sigma) and mixed again at 1,000 rpm for 20 minutes. Following nuclease treatment, samples were clarified by centrifugation at 15,000 × g for 12 minutes. Protein concentrations were determined using the Pierce bicinchoninic acid (BCA) assay (Thermo Fisher Scientific). Disulfide bonds were reduced with 5 mM tris(2-carboxyethyl)phosphine (TCEP, Thermo Fisher Scientific) and alkylated with 10 mM iodoacetamide (IAA, Thermo Fisher Scientific) for 30 minutes at room temperature. Samples were acidified with 25% phosphoric acid to a final concentration of 2.5% and processed using the S-Trap™ (Protifi) protocol. Briefly, six volumes of 50 mM triethylammonium bicarbonate (TEAB, pH 7.2) in methanol were added relative to the total lysate volume, including reducing, alkylating, and acidifying reagents, prior to S-Trap loading. Samples were washed three times with 50 mM TEAB (pH 7.2) in methanol and digested overnight at 37°C using sequencing-grade trypsin and Lys-C (Promega) at a 1:50 enzyme-to-protein ratio in 50 mM TEAB. Peptides were eluted from the S-Trap columns using sequential washes of 50 mM TEAB, 0.2% formic acid (FA), and 50% acetonitrile (ACN) with 0.2% FA. Eluted peptides were quantified using the Pierce Fluorometric Peptide Assay (Thermo Fisher Scientific). For experiments involving paired global proteome analysis, a specified amount of peptide was removed prior to enrichment and desalted separately. The remaining peptides were dried using a vacuum concentrator and resuspended in 100 µL PBS in 0.5 mL Protein LoBind tubes for surface glycopeptide capture.

For glycopeptide capture, streptavidin magnetic beads (Genscript) were added at a ratio of 10 µL per 100 µg of peptide (minimum 5 µL, maximum 50 µL), and samples were incubated for 3 hours at room temperature with mixing at 1,100 rpm. Beads were washed sequentially for 2 minutes each with: 1× PBS, 2× 5 M sodium chloride (NaCl), 2× StimLys buffer (100 mM NaCl, 100 mM glycerol, 50 mM Tris, 1% Triton X-100), 2× 50% ACN, 2× 100 mM sodium carbonate (Na_2_CO₃, pH 11), and 3× 50 mM ABC). Beads were then resuspended in 50 µL of 50 mM ABC and transferred to 1.5 mL or 0.5 mL Protein LoBind tubes. Rapid PNGase F (New England Biolabs), diluted 1:10 in 50 mM ABC, was added at 2 µL per sample, and deglycosylation was carried out overnight at 37°C with mixing at 1,350 rpm for 1.5mL tubes or 1,100 rpm for 0.5mL tubes. The next day, the 50 µL eluate was transferred to styrenedivinylbenzene-reverse phase sulfonate (SDB-RPS) stage tips (Empore) for desalting. Beads were then washed with 50 µL of 50 mM ABC in 20% ACN, and the wash was also transferred to the stage tip, bringing the total load volume to 100 µL. Samples were acidified with 15 µL of 10% trifluoroacetic acid (TFA), bringing the final volume to 115 µL, and centrifuged to bind peptides to the resin. Peptides were washed with 125 µL of 1% TFA in isopropanol (IPA), followed by 125 µL of 0.2% TFA. Peptides were eluted with 80% ACN and 5% ammonium hydroxide (NH_4_OH), collected into LC-MS vials preloaded with 4.5 µL of 0.2% n-dodecyl-β-D-maltoside (DDM), dried, and reconstituted in 9 µL of 3% ACN/1% formic acid (FA). A 4 µL injection was used for liquid chromatography–mass spectrometry (LC-MS) analysis.

Peptide aliquots taken for global proteome analysis were adjusted to 100 µL with 1% FA, and an additional volume of 10% FA was added to bring the final FA concentration to 1% prior to desalting. SDB-RPS stage tips (Empore) were conditioned with 100% ACN, and samples were loaded directly onto the tips. Bound peptides were washed twice with 100 µL of 1% FA in IPA, followed by two washes with 100 µL of 1% FA. Peptides were eluted with 80 µL of 80% ACN containing 5% NH_4_OH, dried by vacuum centrifugation, and reconstituted to a final concentration of 1 µg/µL in 3% ACN with 1% FA. A 1 µL injection (1 µg total) was used for LC-MS/MS analysis.

### WGA-HRP Labeling

For tube-based labeling, cells were transferred to either 0.5 mL or 1.5 mL Protein LoBind tubes (Eppendorf) and washed three times with 300 µL of PBS (pH 6.5) at room temperature. The following volumes correspond to the 0.5 mL format; volumes were scaled proportionally for 1.5 mL tubes and plate-based formats. Cells were incubated with 200 µL of 0.5 µM WGA-HRP (Vector Laboratories) in PBS (pH 6.5) supplemented with 1 mM calcium chloride (CaCl_2_) for 5 minutes on ice. Biotinyl tyramide (Sigma-Aldrich) was added to a final concentration of 0.5 mM by spiking in 2 µL of 50 mM stock in DMSO, followed by thorough mixing. Hydrogen peroxide (30%, Thermo Fisher Scientific) was added to a final concentration of 0.003% (2 µL of 0.3%), and samples were incubated at 37°C for 2 minutes to initiate labeling. Labeling was quenched by adding 200uL of cold quench buffer containing 2 mM sodium pyruvate (Thermo Fisher Scientific), 20 mM sodium ascorbate (Thermo Fisher Scientific), and 10 mM Trolox (Thomas Scientific) in PBS (pH 6.5), followed by two washes with 1X cold quench buffer (1 mM sodium pyruvate, 10 mM sodium ascorbate, and 5 mM Trolox in PBS (pH 6.5)), and one wash with ice-cold PBS (pH 6.5).

For plate-based labeling (6-well plates and 10 cm dishes), the same protocol was applied, with reagent volumes scaled to fully cover the adherent monolayer and all steps performed using manual aspiration. After the final PBS wash, cells were gently scraped into tubes and processed identically to suspension samples. All centrifugation steps were performed at 650 × g for 2.5 minutes, and resulting pellets were frozen at –80°C until lysis.

### WGA-HRP Lysis, Enrichment, and Digestion

Cells were lysed in 1× RIPA buffer (Thermo Fisher Scientific) supplemented with 1% Triton X-100 and 1× Halt Protease Inhibitor Cocktail (100×, Thermo Fisher Scientific). Samples were incubated at 4°C for 30 minutes with gentle mixing at 500 rpm on a thermomixer. Lysates were clarified by centrifugation at 15,000 × g for 12 minutes at 4°C, and protein concentration was determined using the Pierce BCA Protein Assay.

For enrichment, 100 µL of High Capacity NeutrAvidin resin (Thermo Fisher Scientific) was added to each sample and incubated at 4°C for 1 hour with mixing at 1,300 rpm. Samples were transferred to Bio-Rad Bio-Spin columns (Bio-Rad) and washed sequentially with 3 mL each of RIPA buffer, 1 M NaCl in PBS, 50% ACN, 3M Urea in 50mM TEAB, 6M Urea in 50mM TEAB, and 50 mM TEAB. After the final wash, beads were transferred to 1.5 mL Protein LoBind tubes with 200 µL of 50 mM TEAB. Disulfide bonds were reduced with 5 mM TCEP and alkylated with 10 mM IAA for 30 minutes at room temperature. Reagents were removed, and resin was resuspended in 100 µL of 50 mM TEAB containing 1 µg each of trypsin and Lys-C (Promega). Digestion was carried out overnight at room temperature with mixing at 1,300 rpm. The following day, resin was separated using Pierce Snap Cap Spin Columns (Thermo Fisher Scientific), and the peptide-containing flow-through was collected. Peptides were quantified using the Pierce Fluorometric Peptide Assay and acidified to 1% formic acid (FA) using 10% FA.

Peptides were desalted using in-house C18 stage tips prepared with two stacked punches of Empore C18 material (3M). Tips were conditioned sequentially with 100% methanol, 50% ACN containing 0.1% FA, and twice with 0.1% FA. Acidified peptide samples were loaded directly onto the stage tips, followed by two washes with 0.1% FA. Peptides were eluted with 80 µL of 50% ACN containing 0.1% FA and dried by vacuum centrifugation. Final resuspension volumes were adjusted based on peptide yields from the Pierce Fluorometric Peptide Assay to achieve a target concentration of 100 ng/µL in 3% ACN with 1% FA. A total of 2 µL (200 ng) was injected for LC-MS/MS analysis.

### Input Normalization

For HEK293T titration experiments, enrichment was performed on defined cell numbers without protein normalization. For both KMS-12 and A549 samples, input was guided by cell counts or equivalent seeding conditions, and protein amounts were normalized based on BCA measurements to account for minor differences in recovery. This normalization approach was applied consistently across both N-glyco and WGA-HRP workflows.

### LC-MS Analysis

LC-MS/MS analysis was conducted using a Vanquish Neo™ UHPLC system (Thermo Fisher Scientific) coupled to an Orbitrap Exploris™ 480 mass spectrometer (Thermo Fisher Scientific) operating in data-independent acquisition (DIA) mode. Peptides were separated on a 25 cm New Objective PicoFrit™ column (75 µm inner diameter, 10 µm tip ID) packed in-house with 1.5 µm C18 particles (ReproSil-Pur C18-AQ, Dr. Maisch) at a flow rate of 200 nL/min. Surface-enriched samples from both the N-glyco and WGA-HRP workflows were analyzed using an 85-minute active gradient: 1% to 4% buffer B in 1 minute, 4% to 5% in 1 minute, 5% to 24% over 58 minutes, 24% to 28% over 7.5 minutes, 28% to 36% over 7.5 minutes, and 36% to 72% over the final 10 minutes. MS1 scans were acquired from m/z 350–1800 at a resolution of 120,000 with an AGC target of 3E6 and a maximum injection time of 20 ms. MS2 spectra were acquired across m/z 350-1450 using 20 variable windows, at a resolution of 30,000, with an AGC target of 1E6, maximum injection time of 54 ms, and normalized collision energy (NCE) of 27% with a default charge state of +3 (Table S1E).

For global proteome analysis, a 94-minute active gradient was used: 1.6% to 4.8% buffer B in 1 minute, 4.8% to 24% over 84 minutes, and 24% to 48% over the final 9 minutes. MS1 spectra were acquired over m/z 350–1800 with a resolution of 120,000 at 200 m/z, an AGC target of 3E6, and a maximum injection time of 20 ms. DIA was performed using 30 variable windows covering m/z 400-850. MS2 scans were acquired with an AGC target of 5E5, maximum injection time of 54 ms, and NCE set to 27% with a default charge state of +3 (Table S1F).

### Data Analysis

All DIA data were analyzed using Spectronaut (version 19, Biognosys AG) in directDIA mode. Spectra were searched against the UniProt Swiss-Prot human reference proteome (20,421 entries, downloaded June 2025) using Trypsin/P specificity, allowing up to 2 missed cleavages. Carbamidomethylation of cysteine was set as a fixed modification. Oxidation of methionine and protein N-terminal acetylation were included as variable modifications for all datasets. For N-glycoprotein-enriched samples, asparagine deamidation was added as a variable modification to detect PNGase F-cleaved glycosylation sites, and site localization filtering was applied with a minimum probability cutoff of 0.75. Spectronaut’s modification filter was then used to retain only peptides containing deamidation, and these filtered peptides were used for protein quantification. “Exclude De-amidated Peptides” was also turned off in the calibration DIA analysis settings. Only proteins containing at least one peptide mapping to an N-X-S/T consensus motif were retained for downstream analysis. Quantification was performed using Spectronaut’s default protein inference and abundance estimation algorithms.

For WGA-HRP-enriched and global proteome datasets, a standard search workflow was used without the deamidation modification. Protein-level filtering was applied to remove experiment-wide single-peptide identifications. In all cases, protein and precursor identifications were filtered at 1% FDR using Spectronaut’s q-value score.

For CSP ID figures (HEK titration, KMS12 and A549 high/low input, endometrial titration, and endometrial method evaluation), the method evaluation option in Spectronaut was enabled to minimize match-between-runs (MBR) effects on identification rates and obtain more accurate identifications reflective of each input amount or dissociation technique. All figures referencing quantitative changes or comparisons were generated without method evaluation enabled, per Spectronaut’s recommendations for accurate quantification.

Protein intensity values were log_2_-transformed and normalized using sample-wise median centering in R (Protigy pipeline), to account for global differences in sample loading and MS signal. For the HEK titration and KMS12 fresh/frozen experiments, normalization was performed within each group, not experiment-wide. For comparative analyses, fold changes between conditions were calculated based on log_2_-transformed protein intensities. For differential abundance analysis, two-sample *t*-tests were performed, and results were visualized as volcano plots displaying log_2_ fold-change versus –log₁₀ *p*-value (nominal *p*-values reported). All statistical analyses were performed in R or Python, and figures were generated using ggplot2, matplotlib, or GraphPad Prism as appropriate.

---

**Figure S1.**
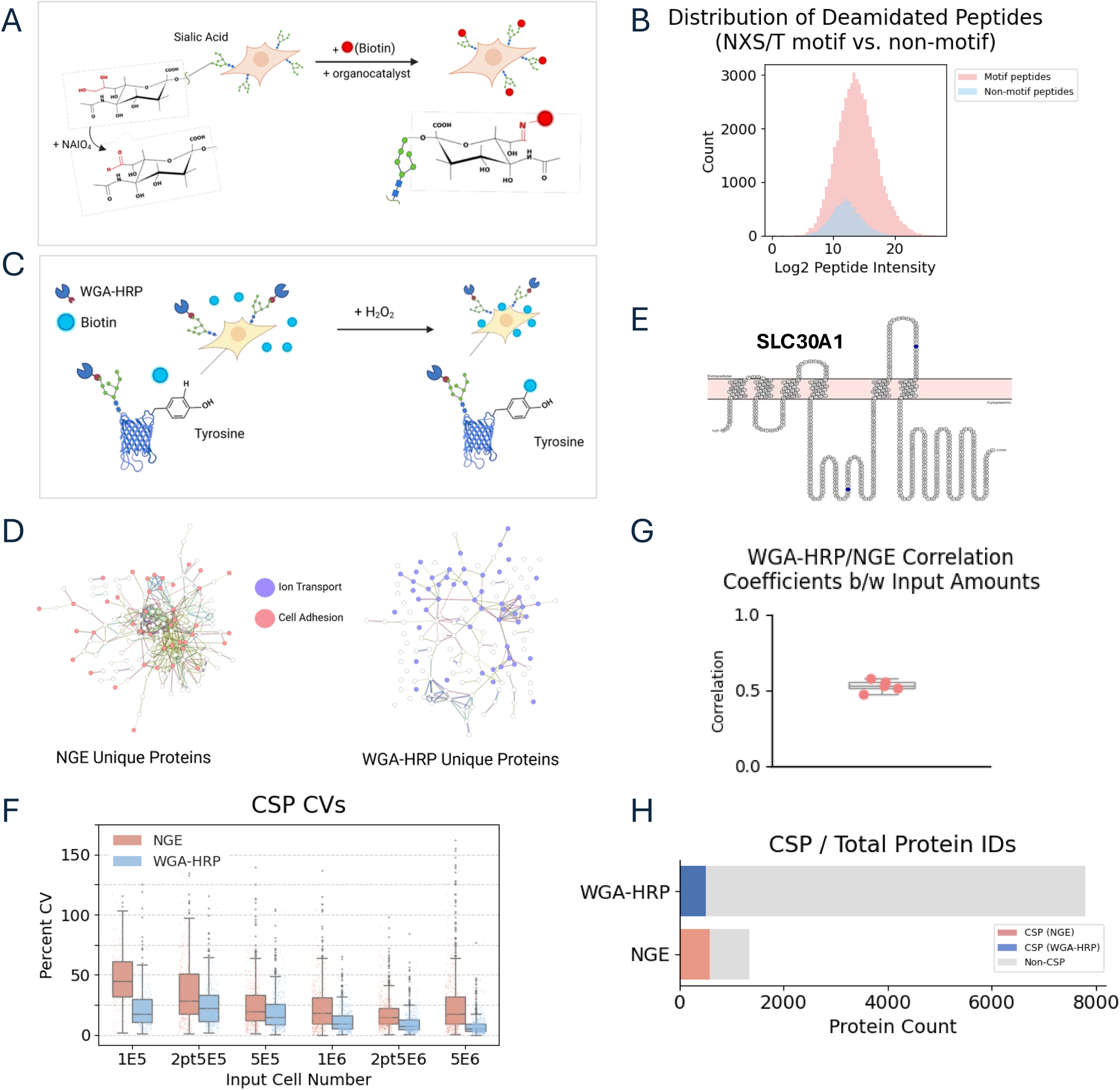
Comparative strategies for cell surface protein labeling and additional HEK titration metrics. **A.** Cell surface glycan labeling by NaIO_4_ oxidation and amine-based biotinylation. Sialic acid residues are selectively oxidized to aldehydes and reacted with an amine-containing biotin reagent in the presence of an organocatalyst, 5-methoxyanthranilic acid (5MA). **B.** Log_2_-transformed peptide intensity distributions for motif-containing and non-motif peptides from N-glyco–enriched samples (across all HEK293T inputs, includes overlapping peptides between inputs) were classified based on confident deamidation site localization (localization probability ≥ 0.75) and N-X-S/T motif presence, with motif-containing proteins retained for downstream CSP analysis. **C.** Cell surface protein labeling by WGA-HRP and biotin-tyramide. WGA-HRP binds to terminal GlcNAc and sialic acids, enabling HRP-catalyzed deposition of biotin-tyramide onto proximal tyrosine residues. **D.** STRING network analysis of proteins uniquely identified by each method, annotated for functions in cell adhesion and ion transport with BH FDR < 0.01. **E.** Representative topology of the ion transport protein SLC30A1, uniquely identified by WGA-HRP. The diagram was generated using the Protter web-based application, with blue dots indicating N-X-S/T motifs. **F.** Coefficient of variation (CV) percentages for fully quantified proteins across replicates at each input amount, assessing quantitative reproducibility (N = 3). **G.** Pearson correlation coefficients between glycan oxidation– and WGA-HRP–based surfaceome data across input amounts (1×10^5^ to 5×10^6^ cells). **H.** Cell surface protein identifications compared to total number of proteins cumulatively across all HEK titration input amounts for both protocols.

**Figure S2.**
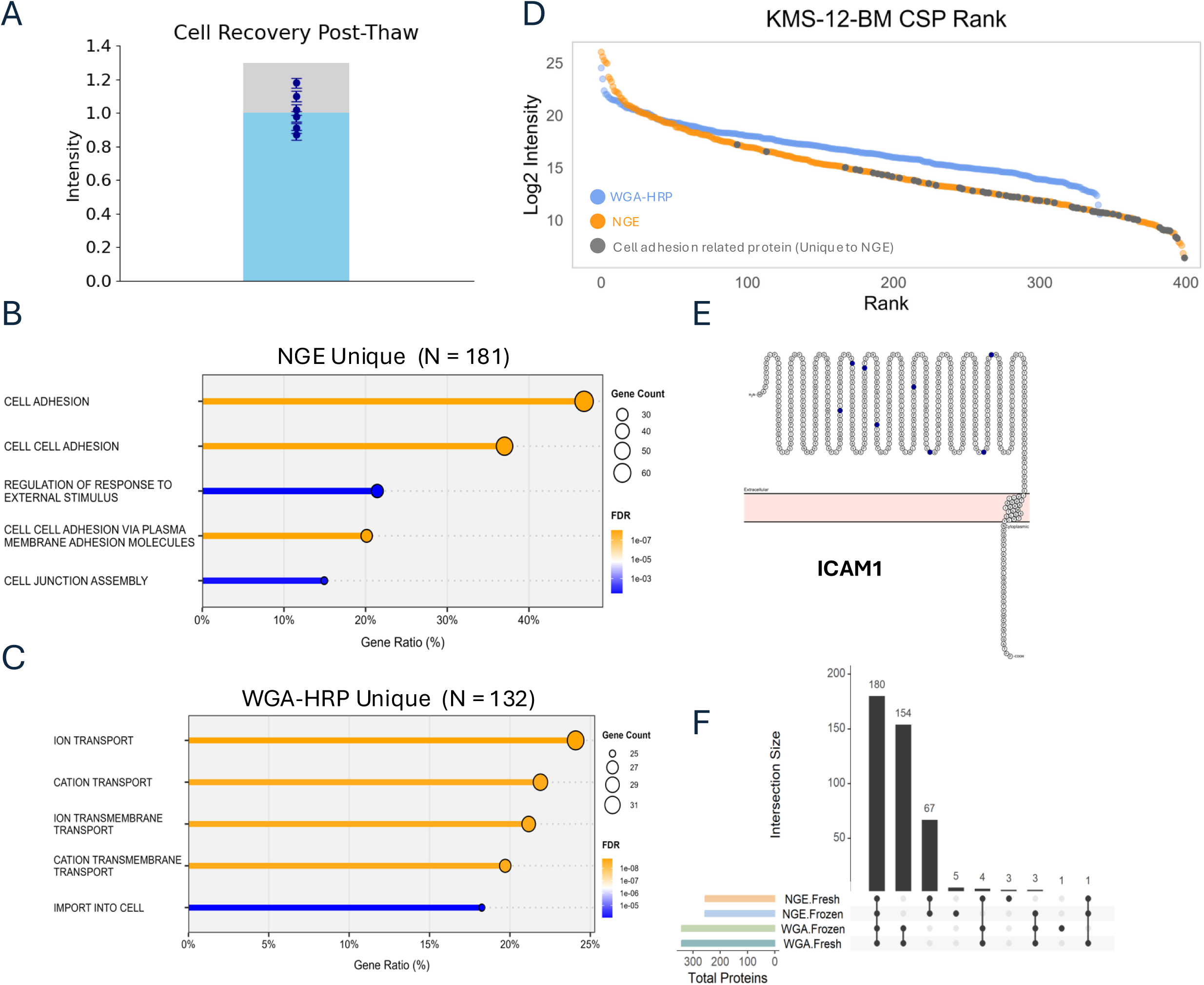
Method-specific surface proteome features and agreement metrics for KMS-12-BM suspension cell line. **A.** Bar plot of post-thaw cell recovery, with pre-freeze cell count (1.3 × 10^6^) shown as a light gray background bar. Average cell recovery was 1.1 × 10^6^ (N = 6). **B–C.** GO Biological Process overrepresentation analysis for proteins uniquely identified by each method. Enriched terms for NGE-unique proteins are dominated by cell adhesion processes; ion transport processes are enriched for WGA-HRP–unique proteins. **D.** Rank plots showing the dynamic range of surface protein intensities for each enrichment method in KMS-12-BM cells. **E.** Representative topological map of a cell adhesion protein identified by N-glyco enrichment (ICAM1), highlighting a large extracellular domain with multiple predicted N -glycosylation motifs. The diagram was generated using the Protter web-based application, with blue dots indicating N-X-S/T motifs. **F.** Upset plot showing the number of intersecting proteins among fresh and frozen samples labeled with either NGE or WGA-HRP. Horizontal bars represent total number of proteins identified in each dataset.

**Figure S3.**
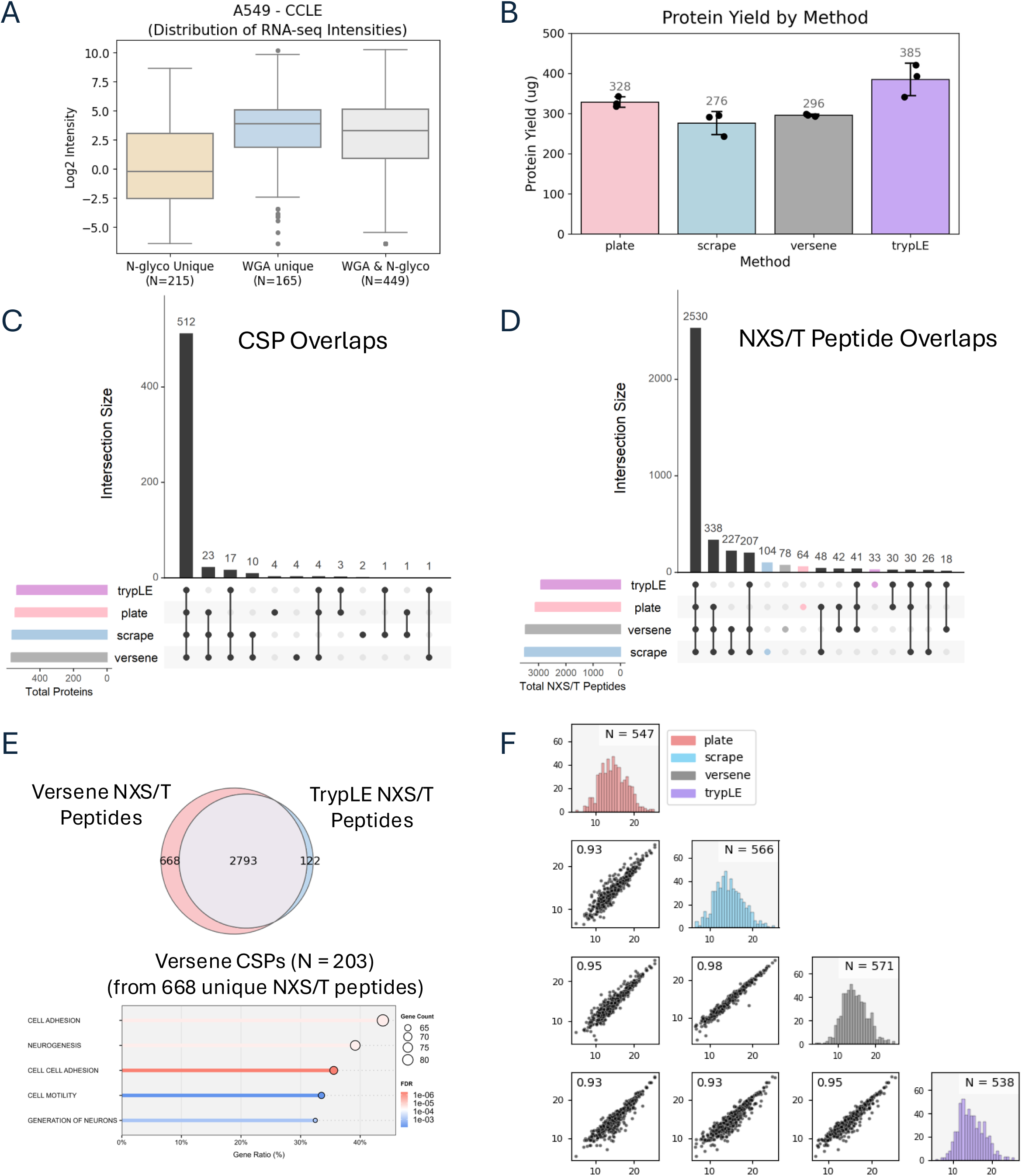
Method-specific surface proteome features and agreement metrics for A549 adherent cell line. **A.** RNA expression levels (log_2_ intensity) for surface proteins uniquely identified by N-glyco enrichment (N = 215), WGA enrichment (N = 165), or detected by both methods (N = 449) in A549 cells, using CCLE RNA-seq data. **B.** Total protein yield (µg) across four dissociation methods (plate, scrape, versene, trypsin-based TrypLE) from equal cell inputs. Each dot represents an independent replicate. **C.** Overlap of cell surface proteins (CSPs) identified across the four dissociation methods. The largest shared set (N = 516) is common to all methods, with smaller method-specific subsets. **D.** Overlap of identified NXS/T-containing peptides across the four dissociation methods. **E.** Gene Ontology Biological Process (GO:BP) over-representation analysis of CSPs detected from the versene condition (N = 210) within the 696 unique NXS/T peptides. Top enriched terms include neurogenesis, cell adhesion, and motility. **F.** Scatter plots and histograms showing consistent surface protein quantification across A549 6-well plate conditions (plate-based labeling and various cell detachments) for the NGE method.

**Figure S4.**
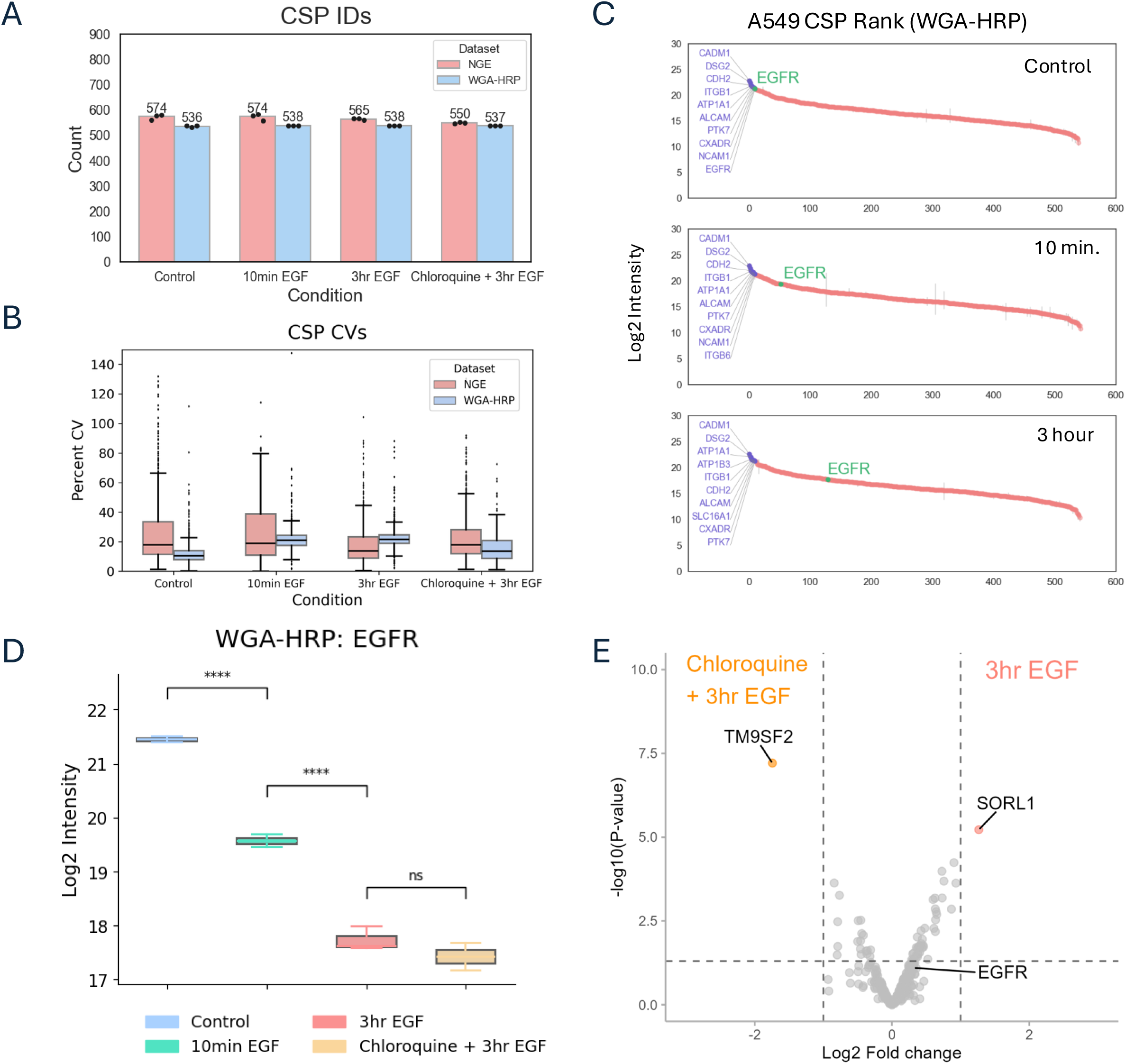
Method-specific surfaceome performance and EGFR detection across receptor trafficking conditions. **A.** Surface protein identifications across the four treatment conditions from NGE and WGA-HRP–based surfaceome profiling (N = 3). **B.** Coefficients of variation (CVs) across the four treatment conditions from NGE and WGA -HRP–based surfaceome profiling (N = 3). **C.** Rank plot of WGA-HRP–based surfaceome data from control, 10 min EGF, and 3 hr EGF conditions, highlighting EGFR and high-abundance stable surface proteins. **D.** EGFR abundance measured by WGA-HRP across the four conditions, showing consistency with NGE-based trends (N=3). Significance determined by unpaired two-tailed t-tests: *P* > 0.05 (NS), *P* < 0.0001 (****). **E.** Volcano plot of WGA-HRP surfaceome data comparing 3-hour EGF vs. chloroquine + 3-hour EGF, showing downregulation of EGFR recycling proteins in chloroquine-pretreated samples, with less pronounced EGFR suppression—potentially due to intracellular EGFR contributing to background signal (log_2_-fold change > |1|; p < 0.05 by t-test).

**Figure S5.**
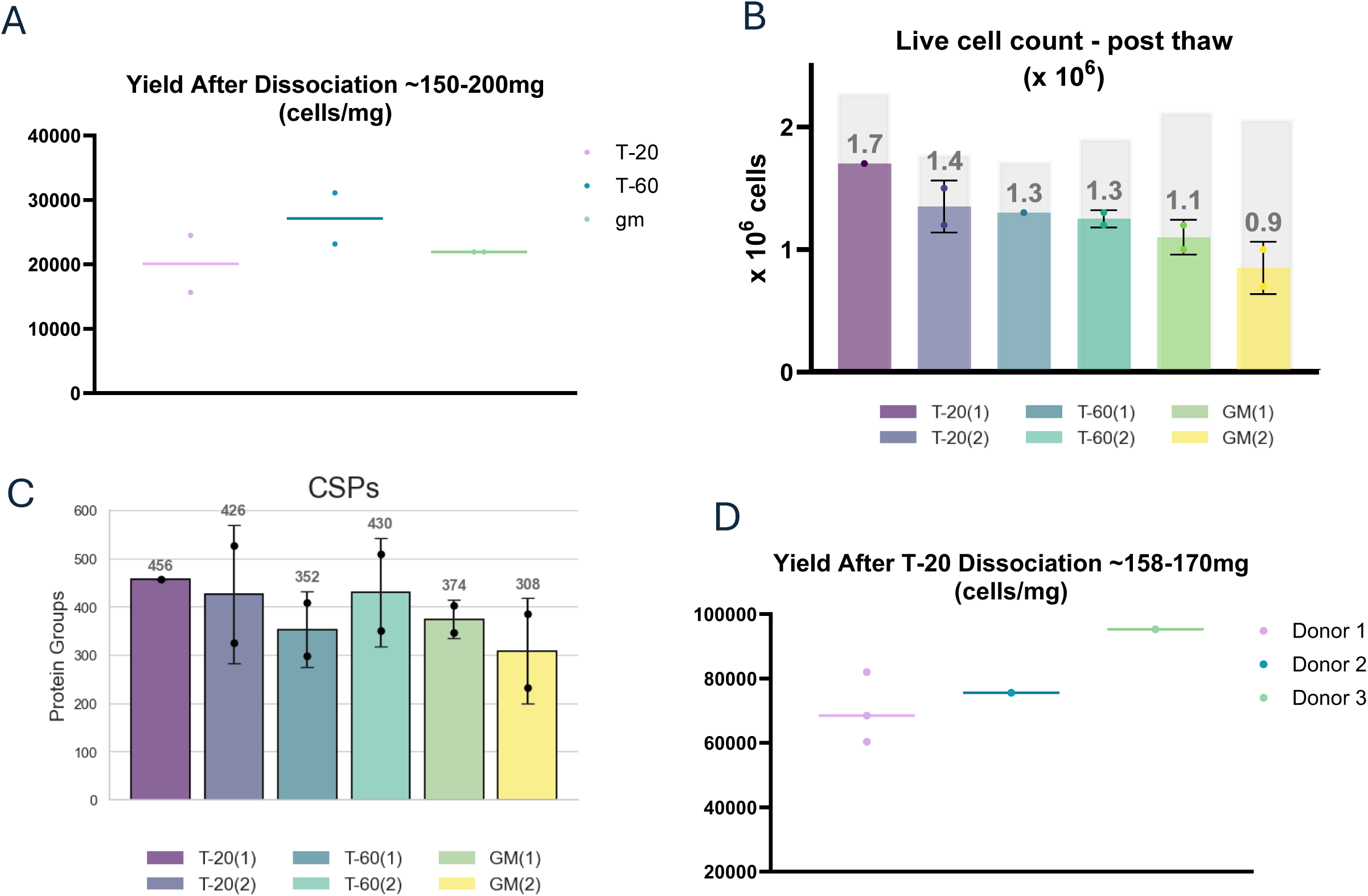
Evaluation of tissue dissociation strategies for cryopreserved surface proteomics. **A.** Workflow for dissociation and cryopreservation of fresh endometrial tissue using three protocols: T-20, T-60, and gentleMACS (GM). **B.** Post-thaw live cell counts following cryopreservation. Bars show the number of viable cells recovered after thawing, based on two biological replicates per condition (e.g., T-20(1), T-20(2)), each derived from independent tissue pieces. Error bars represent replicate cryopreserved vials from the same dissociation, processed and sorted with matched cell numbers. Light gray bars indicate live cell counts prior to cryopreservation. **C.** Number of confident cell surface proteins (CSPs) identified across all cryopreserved samples. Each bar represents a biological replicate; error bars show technical replicate variability following surface enrichment and MS analysis. **D.** Cell yields obtained during T-20 dissociation of three independent donor samples (158–170 mg tissue per sample), used in downstream titration experiments. Cell recovery ranged from ∼60,000 to 90,000 cells per mg of tissue.

## REFERENCES

1. Bausch-Fluck, D., Hofmann, A., Bock, T., Frei, A. P., Cerciello, F., Jacobs, A., Moest, H., Omasits, U., Gundry, R. L., Yoon, C., Schiess, R., Schmidt, A., Mirkowska, P., Härtlová, A., Van Eyk, J. E., Bourquin, J.-P., Aebersold, R., Boheler, K. R., Zandstra, P., and Wollscheid, B. (2015) A mass spectrometric-derived cell surface protein atlas. PLoS One 10, e0121314

2. Wollscheid, B., Bausch-Fluck, D., Henderson, C., O’Brien, R., Bibel, M., Schiess, R., Aebersold, R., and Watts, J. D. (2009) Erratum: Corrigendum: Mass-spectrometric identification and relative quantification of N-linked cell surface glycoproteins. Nat. Biotechnol. 27, 864–864

3. Jang, J. H., and Hanash, S. (2003) Profiling of the cell surface proteome. Proteomics 3, 1947–1954

4. Udeshi, N. D., Xu, C., Jiang, Z., Gao, S. M., Yin, Q., Luo, W., Carr, S. A., Davis, M. M., and Li, J. (2024) Cell-surface milieu remodeling in human dendritic cell activation. J. Immunol. 213, 1023–1032

5. Wollscheid, B., Bausch-Fluck, D., Henderson, C., O’Brien, R., Bibel, M., Schiess, R., Aebersold, R., and Watts, J. D. (2009) Mass-spectrometric identification and relative quantification of N-linked cell surface glycoproteins. Nat. Biotechnol. 27, 378–386

6. Ferguson, I. D., Patiño-Escobar, B., Tuomivaara, S. T., Lin, Y.-H. T., Nix, M. A., Leung, K. K., Kasap, C., Ramos, E., Nieves Vasquez, W., Talbot, A., Hale, M., Naik, A., Kishishita, A., Choudhry, P., Lopez-Girona, A., Miao, W., Wong, S. W., Wolf, J. L., Martin, T. G., 3rd, Shah, N., Vandenberg, S., Prakash, S., Besse, L., Driessen, C., Posey, A. D., Jr, Mullins, R. D., Eyquem, J., Wells, J. A., and Wiita, A. P. (2022) The surfaceome of multiple myeloma cells suggests potential immunotherapeutic strategies and protein markers of drug resistance. Nat. Commun. 13, 4121

7. Huang, P., Gao, W., Fu, C., Wang, M., Li, Y., Chu, B., He, A., Li, Y., Deng, X., Zhang, Y., Kong, Q., Yuan, J., Wang, H., Shi, Y., Gao, D., Qin, R., Hunter, T., and Tian, R. (2025) Clinical functional proteomics of intercellular signalling in pancreatic cancer. Nature 637, 726–735

8. Gundry, R. L., Riordon, D. R., Tarasova, Y., Chuppa, S., Bhattacharya, S., Juhasz, O., Wiedemeier, O., Milanovich, S., Noto, F. K., Tchernyshyov, I., Raginski, K., Bausch-Fluck, D., Tae, H.-J., Marshall, S., Duncan, S. A., Wollscheid, B., Wersto, R. P., Rao, S., Van Eyk, J. E., and Boheler, K. R. (2012) A cell surfaceome map for immunophenotyping and sorting pluripotent stem cells. Mol. Cell. Proteomics 11, 303–316

9. Dieters-Castator, D. Z., Manzanillo, P., Yang, H.-Y., Modak, R. V., Rardin, M. J., and Gibson, B. W. (2024) Magnetic bead-based workflow for sensitive and streamlined cell surface proteomics. J. Proteome Res. 23, 618–632

10. Kirkemo, L. L., Elledge, S. K., Yang, J., Byrnes, J. R., Glasgow, J. E., Blelloch, R., and Wells, J. A. (2022) Cell-surface tethered promiscuous biotinylators enable comparative small-scale surface proteomic analysis of human extracellular vesicles and cells. Elife 11,

11. Vilen, Z., Reeves, A. E., O’Leary, T. R., Joeh, E., Kamasawa, N., and Huang, M. L. (2023) Cell surface engineering enables surfaceome profiling. ACS Chem. Biol. 18, 701–710

12. Li, J., Han, S., Li, H., Udeshi, N. D., Svinkina, T., Mani, D. R., Xu, C., Guajardo, R., Xie, Q., Li, T., Luginbuhl, D. J., Wu, B., McLaughlin, C. N., Xie, A., Kaewsapsak, P., Quake, S. R., Carr, S. A., Ting, A. Y., and Luo, L. (2020) Cell-surface proteomic profiling in the fly brain uncovers wiring regulators. Cell 180, 373–386.e15

13. Shuster, S. A., Li, J., Chon, U., Sinantha-Hu, M. C., Luginbuhl, D. J., Udeshi, N. D., Carey, D. K., Takeo, Y. H., Xie, Q., Xu, C., Mani, D. R., Han, S., Ting, A. Y., Carr, S. A., and Luo, L. (2022) In situ cell-type-specific cell-surface proteomic profiling in mice. Neuron 110, 3882–3896.e9

14. Luecke, L. B., Waas, M., Littrell, J., Wojtkiewicz, M., Castro, C., Burkovetskaya, M., Schuette, E. N., Buchberger, A. R., Churko, J. M., Chalise, U., Waknitz, M., Konfrst, S., Teuben, R., Morrissette-McAlmon, J., Mahr, C., Anderson, D. R., Boheler, K. R., and Gundry, R. L. (2023) Surfaceome mapping of primary human heart cells with CellSurfer uncovers cardiomyocyte surface protein LSMEM2 and proteome dynamics in failing hearts. Nat. Cardiovasc. Res. 2, 76–95

15. Donlin, L. T., Accelerating Medicines Partnership RA/SLE Network, Rao, D. A., Wei, K., Slowikowski, K., McGeachy, M. J., Turner, J. D., Meednu, N., Mizoguchi, F., Gutierrez-Arcelus, M., Lieb, D. J., Keegan, J., Muskat, K., Hillman, J., Rozo, C., Ricker, E., Eisenhaure, T. M., Li, S., Browne, E. P., Chicoine, A., Sutherby, D., Noma, A., Nusbaum, C., Kelly, S., Pernis, A. B., Ivashkiv, L. B., Goodman, S. M., Robinson, W. H., Utz, P. J., Lederer, J. A., Gravallese, E. M., Boyce, B. F., Hacohen, N., Pitzalis, C., Gregersen, P. K., Firestein, G. S., Raychaudhuri, S., Moreland, L. W., Holers, V. M., Bykerk, V. P., Filer, A., Boyle, D. L., Brenner, M. B., and Anolik, J. H. (2018) Methods for high-dimensional analysis of cells dissociated from cryopreserved synovial tissue. Arthritis Res. Ther. 20,

16. Wu, S. Z., Roden, D. L., Al-Eryani, G., Bartonicek, N., Harvey, K., Cazet, A. S., Chan, C.-L., Junankar, S., Hui, M. N., Millar, E. A., Beretov, J., Horvath, L., Joshua, A. M., Stricker, P., Wilmott, J. S., Quek, C., Long, G. V., Scolyer, R. A., Yeung, B. Z., Segara, D., Mak, C., Warrier, S., Powell, J. E., O’Toole, S., Lim, E., and Swarbrick, A. (2021) Cryopreservation of human cancers conserves tumour heterogeneity for single-cell multi-omics analysis. Genome Med. 13, 81

17. van Oostrum, M., Müller, M., Klein, F., Bruderer, R., Zhang, H., Pedrioli, P. G. A., Reiter, L., Tsapogas, P., Rolink, A., and Wollscheid, B. (2019) Classification of mouse B cell types using surfaceome proteotype maps. Nat. Commun. 10, 5734

18. Fonseca, M. A. S., Haro, M., Wright, K. N., Lin, X., Abbasi, F., Sun, J., Hernandez, L., Orr, N. L., Hong, J., Choi-Kuaea, Y., Maluf, H. M., Balzer, B. L., Fishburn, A., Hickey, R., Cass, I., Goodridge, H. S., Truong, M., Wang, Y., Pisarska, M. D., Dinh, H. Q., El-Naggar, A., Huntsman, D. G., Anglesio, M. S., Goodman, M. T., Medeiros, F., Siedhoff, M., and Lawrenson, K. (2023) Single-cell transcriptomic analysis of endometriosis. Nat. Genet. 55, 255–267

19. Gołąbek-Grenda, A., and Olejnik, A. (2022) In vitro modeling of endometriosis and endometriotic microenvironment - Challenges and recent advances. Cell. Signal. 97, 110375

20. Hung, V., Udeshi, N. D., Lam, S. S., Loh, K. H., Cox, K. J., Pedram, K., Carr, S. A., and Ting, A. Y. (2016) Spatially resolved proteomic mapping in living cells with the engineered peroxidase APEX2. Nat. Protoc. 11, 456–475

21. Rhee, H.-W., Zou, P., Udeshi, N. D., Martell, J. D., Mootha, V. K., Carr, S. A., and Ting, A. Y. (2013) Proteomic mapping of mitochondria in living cells via spatially restricted enzymatic tagging. Science 339, 1328–1331

22. Bausch-Fluck, D., Goldmann, U., Müller, S., van Oostrum, M., Müller, M., Schubert, O. T., and Wollscheid, B. (2018) The in silico human surfaceome. Proc. Natl. Acad. Sci. U. S. A. 115, E10988–E10997

23. Hallgren, J., Tsirigos, K. D., Pedersen, M. D., Almagro Armenteros, J. J., Marcatili, P., Nielsen, H., Krogh, A., and Winther, O. (2022) DeepTMHMM predicts alpha and beta transmembrane proteins using deep neural networks. bioRxiv,

24. Omasits, U., Ahrens, C. H., Müller, S., and Wollscheid, B. (2014) Protter: interactive protein feature visualization and integration with experimental proteomic data. Bioinformatics 30, 884–886

25. Henriksen, L., Grandal, M. V., Knudsen, S. L. J., van Deurs, B., and Grøvdal, L. M. (2013) Internalization mechanisms of the epidermal growth factor receptor after activation with different ligands. PLoS One 8, e58148

26. Sigismund, S., Argenzio, E., Tosoni, D., Cavallaro, E., Polo, S., and Di Fiore, P. P. (2008) Clathrin-mediated internalization is essential for sustained EGFR signaling but dispensable for degradation. Dev. Cell 15, 209–219

27. Liu, L., Han, C., Yu, H., Zhu, W., Cui, H., Zheng, L., Zhang, C., and Yue, L. (2018) Chloroquine inhibits cell growth in human A549 lung cancer cells by blocking autophagy and inducing mitochondrial-mediated apoptosis. Oncol. Rep.,

28. Jiang, Z., Bi, F., Ge, Z., Mansolf, M., Hartwich, T. M. P., Kolesnyk, V., Yang, K., Park, W., Kim, D., Grechukhina, O., Hui, P., Kim, S. W., and Yang-Hartwich, Y. (2025) SORL1-mediated EGFR and FGFR4 regulation enhances chemoresistance in ovarian cancer. Cancers (Basel) 17,

29. Pietilä, M., Sahgal, P., Peuhu, E., Jäntti, N. Z., Paatero, I., Närvä, E., Al-Akhrass, H., Lilja, J., Georgiadou, M., Andersen, O. M., Padzik, A., Sihto, H., Joensuu, H., Blomqvist, M., Saarinen, I., Boström, P. J., Taimen, P., and Ivaska, J. (2019) SORLA regulates endosomal trafficking and oncogenic fitness of HER2. Nat. Commun. 10, 2340

